# Genetic divergence in the absence of strong ecological differences between coexisting white and common Atlantic marine sticklebacks

**DOI:** 10.1101/2025.09.30.678994

**Authors:** Kieran Samuk, Hannah Visty, Dolph Schluter

## Abstract

Identifying taxa in the earliest phases of speciation is critical for understanding how reproductive isolation arises. In Nova Scotia, Canada, “white” threespine sticklebacks co-occur with common marine sticklebacks but differ in nuptial coloration, nesting behavior, and parental care, raising the possibility that they represent a nascent species. We combined population genomics, morphometrics, and stable isotope analysis to test whether white sticklebacks represent a distinct lineage and whether they have diverged along ecological axes as in other stickleback populations. Genotyping-by-sequencing revealed that male and female white sticklebacks form a genetic cluster distinct from sympatric common sticklebacks with evidence of ongoing gene flow yet with very low overall genomic divergence (F_ST_ ≈ 0.01). Genetic differences were distributed across many loci rather than localized to a single genomic region. Morphological and isotopic analyses revealed no differences in most classic ecological traits (body shape, armor, gill rakers, or trophic niche). Instead, whites are smaller-bodied, paler, and exhibit shorter spines, reduced testes size, and smaller but more numerous eggs compared to common sticklebacks. These results indicate that white sticklebacks are genetically distinct from the common Atlantic threespine stickleback but have not diverged conspicuously in their ecology, suggesting that their differentiation is driven by reproductive and sexual traits rather than trophic specialization. The white stickleback thus represents a promising new system for investigating the interplay of sexual selection, reproductive strategy, and gene flow in the early stages of speciation.

## Introduction

Understanding the genetic and phenotypic changes that lead to the formation of new species remains a major goal of evolutionary biology (Coyne & Orr, 2004; Knott et al., 2012). While recent advances in genomics have provided insight into some aspects of the evolution of reproductive isolation (RI), there are still many unanswered questions about how isolation evolves in natural systems (R. Butlin et al., 2012; Noor & Feder, 2006; Wu & Ting, 2004). How do new species arise in the face of gene flow? What role does divergent natural selection play in the formation of species boundaries? What phenotypic and genetic changes initiate the speciation process?

Choosing a study system to approach these questions is complicated by the fact that species vary in their progress along the “speciation continuum” (Roux et al., 2016). It is generally agreed that younger species are the most useful for studying speciation (Coyne & Orr, 2004; Knott et al., 2012; Via, 2009). This is because recently diverged species avoid a key problem with the study of the evolution of reproductive isolation (hereafter, RI): as diverging populations proceed toward reproductive isolation, new reproductive barriers arise and mask those that formed at the onset of speciation (R. Butlin et al., 2012; Coyne & Orr, 2004; Roux et al., 2016; Via, 2009). These later-forming barriers may help maintain RI (e.g. by causing post-zygotic RI), but they are not necessarily informative of the key barriers that originally caused speciation to occur (Coyne & Orr, 2004; Orr, 2005; Price, 2008). For example, a late-evolved lethal intrinsic incompatibility between two species could mask the role of poor ecological performance of hybrids because hybrids are never formed (Butlin et al., 2014). Later-accumulating barriers also attenuate gene flow and cause genome-wide divergence to increase, reducing the power of divergence-based methods for detecting loci involved in RI (Egan et al., 2013; Feder et al., 2012; Noor & Feder, 2006). Thus, we can maximize our ability to find the genetic and phenotypic changes that initiate speciation by studying recently diverged taxa, i.e. young species.

The utility of young co-occurring species for identifying RI has become particularly apparent in the age of genomics (Feder et al., 2012; Ravinet et al., 2017). This is because young species often exhibit partial reproductive isolation, which allows gene flow to homogenize parts of the genome that are not involved in the maintenance of species differences, amplifying the genomic signatures of divergence at RI loci (Rogers & Bernatchez, 2006; Stephan et al., 2010; Wu, 2001). Interestingly, despite these obvious advantages, there are still only a handful of developed systems for studying the very early stages of speciation, particularly in the presence of gene flow. Some examples of such systems include *Rhagoletus* apple/hawthorn flies, *Littorina* intertidal snails, *Helianthus* dune sunflowers and *Timema* walking sticks, which have all begun to yield key insights into the speciation process (Feder et al., 2003; Le Moan et al., 2023; Nosil et al., 2005; Ostevik et al., 2016; Rieseberg et al., 2012). A complete picture of the early phases of speciation will require additional study systems, particularly those with developed genomic resources.

The threespine stickleback (*Gasterosteus aculeatus*) species complex is thought to harbor many young species. In five post-glacial lake systems, stickleback species pairs are genetically diverged and exhibit strong but incomplete RI (Foster & Bell, 1994; Hendry et al., 2013; McKinnon & Rundle, 2002; Reid et al., 2021). Gene flow has apparently occurred throughout divergence (Wang, 2018), and RI is mediated largely by their ecological differences (e.g. selection against hybrids with mismatched trophic traits; Arnegard et al 2014, Schluter et al 2025) and by assortative mating on the basis of body size and shape (Conte & Schluter, 2013; Rundle et al., 2005; Bay et al 2017). This pattern of divergence along ecological axes is consistent with the idea that speciation with gene flow can result from strong, divergent natural selection (e.g. provided by different trophic niches). However, many of the best-studied population and species pairs have moderate to high levels of genome-wide genetic differentiation (Hohenlohe et al., 2010; Jones et al., 2012; Reid et al., 2021; Roesti et al., 2012). Thus, we have an incomplete sample of divergence/speciation continuum in sticklebacks and are limited in our ability to probe the genetic and phenotypic changes that underlie the crucial initial stages of speciation. To amend this, new systems are needed in which stickleback species have evolved recently and still exchange genes.

### The white stickleback

The “white” threespine stickleback from Nova Scotia, Canada, may be one such system (Blouw & Hagen, 1990). White sticklebacks appear to be distinct from common marine sticklebacks (hereafter “common sticklebacks”), and both types are broadly sympatric in marine and estuarine environments in Nova Scotia (Blouw & Hagen, 1990). Male white stickleback build nests near the shore (sometimes in the intertidal), and use filamentous algae rather than sand and gravel as nesting substrate (Jamieson et al., 1992a, 1992b; Macdonald et al., 1995). When on the breeding grounds, male white stickleback exhibit striking pearlescent-white breeding colors, instead of the more common olive/blue colors (Blouw & Hagen, 1990). Intriguingly, male white stickleback also lack the classic paternal care behaviors characteristic of male common stickleback: instead of caring for eggs after fertilization, white males carry eggs away from their nest (often out of their territory entirely), disperse them into the surrounding algae, and return to soliciting matings from females (Blouw, 1996; Jamieson et al., 1992b). Male white sticklebacks are also on average ∼20% shorter in body length than common male sticklebacks, resulting in a bimodal distribution of male body sizes at sites where both are found (Blouw & Hagen, 1990).

Recent work has explored a wide variety of phenotypic and genetic differences between white and common sticklebacks. The difference in male nuptial coloration appears to be caused by a reduced density and extent of melanophores in the skin of white males (Haley et al., 2019). Parenting white and common males also differ in total gene expression during the courtship and parental phases (Barbasch et al., 2024), which appears to be underlain by changes in expression in specific regions of the brain (Dan et al., 2024). Differences in parental care behavior have also now been mapped to specific QTL (Behrens, Tucker, et al., 2025), implying a genetic basis. Finally, lab-reared F1 and F2 hybrids between white and common sticklebacks also exhibit transgressive and putatively maladaptive parental care behavior, including reduced fanning behavior, a mix of common and white parental care strategies (e.g. nest building with no fanning and vice versa), and increased rates of egg cannibalism (Behrens, Maciejewski, et al., 2025).

While our understanding of this system is growing, it remains unclear whether white and common sticklebacks represent distinct species. White and common sticklebacks are fully interfertile in advanced generation laboratory crosses (Behrens, Tucker, et al., 2025; Blouw, 1996), suggesting little intrinsic postzygotic isolation. An allozyme study found no evidence of genetic differentiation between the two types (Haglund et al., 1990). However, a distinct class of small-bodied females is always found at sites with small-bodied male white sticklebacks (Blouw & Hagen, 1990; Jamieson et al., 1992a). Mate-choice experiments and field observations suggest that these small females and males mate assortatively (Jamieson et al., 1992b, 1992a). This is consistent with experimental evidence of the role of body size in mate choice in other threespine stickleback (Conte & Schluter, 2013), and suggests there may be pre-mating barriers to gene flow in this system.

These findings suggests that white stickleback may be a fruitful system for studying recent and ongoing speciation. However, assessing the stage of speciation (recent or old), as well as the presence of on-going gene flow requires a detailed study of the genetic relationship between the white and common sticklebacks using modern tools. Here, we employ genetic, morphological, and isotopic data explore the evolutionary history of the white stickleback. We sought the answer the following two main questions:

First, do white sticklebacks represent a nascent species or a complex intrapopulation polymorphism? To answer this question, we examined patterns of genetic polymorphism in wild-caught samples of white and common sticklebacks. If white and common sticklebacks represent separate, distinct lineages in broad sympatry, we expect divergent at sites across the genome rather than at a single site such as is seen in social and reproductive polymorphisms (e.g. Küpper et al. 2016).

Second, have white and common sticklebacks diverged along a trophic ecological axis, as in the limnetic-benthic species pairs? To answer this question, we examined morphological traits and stable isotopes known to be associated with differences in diet and habitat in other threespine stickleback. If white and common sticklebacks have, like other stickleback species pairs, diverged along a trophic axis, we would expect them to display significant differences in ecomorphological traits and isotopic ratios.

By answering these questions, we hope to better understand the early stages of speciation in sticklebacks and lay the groundwork for developing the white stickleback into a full-fledged study system.

## Methods

### Sample collection

In early May-July of 2012 and 2014, we collected white and common threespine stickleback at 16 sites in Nova Scotia, Canada (Table S1). Locations included 11 sites adjacent to the mainland of Nova Scotia and five sites within Bras d’Or Lake, an inland sea on Cape Breton Island. We determined sites using the list of sites in Blouw et al. (1992) as a guide. We focused on sites where both types were most likely to co-occur during the breeding season according to Blouw’s environmental analysis: brackish water with abundant filamentous algae. This sampling scheme ultimately resulted in examining every accessible freshwater estuary we could access by car or short hike along the southern coast of Nova Scotia, from Yarmouth and through the Strait of Canso to Antigonish. In Bras d’Or Lake (2014 only), we sampled estuarine sites (where rivers or other freshwater bodies mixed into the sea) that we could access with a radius of approximately 100km centered on the town of Whycocomagh, NS (Figure 1C).

**Figure 1.**
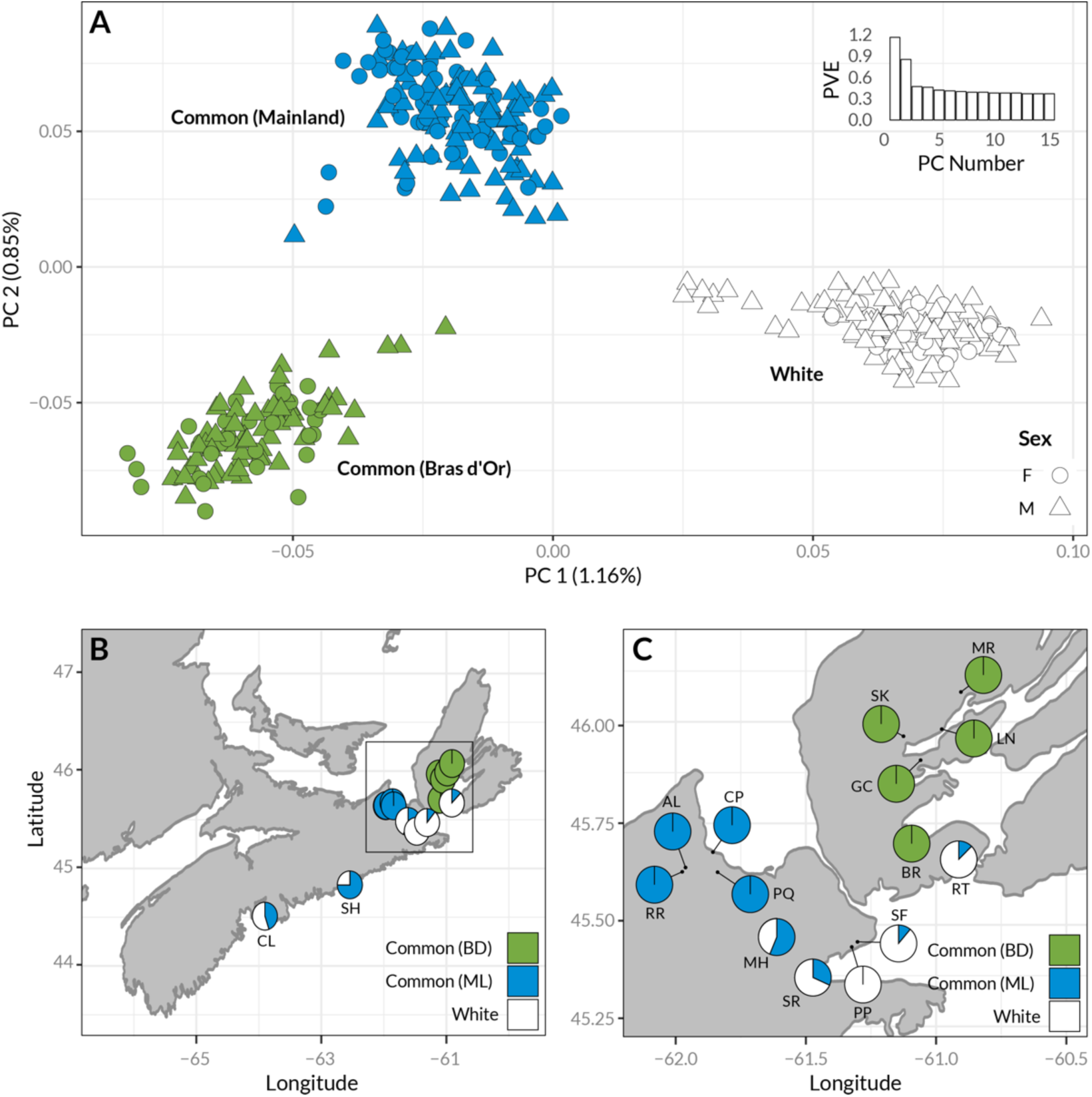
Male and female white sticklebacks form a distinct genotypic cluster. (A) Principal component analysis of ∼8000 LD-pruned SNPs derived from GBS reads. Colors indicate K-means cluster groupings (K=3). (B-C) Maps of the geographic distribution of genotypic clusters in Nova Scotia, Canada. Pie chart sections represent the proportion individuals at each site belonging to each genotypic cluster in A. Site labels: CL = Canal Lake, SH = Sheet Harbour, RR = Rights River, AL = Antigonish Landing, CP = Captain’s Pond, PQ = Pomquet, MH = Milford Haven River, SF = St. Francis Harbour, PP = Porper Pond, SR = Salmon River, RT = River Tillard, BR = Black River, GC = Gillies Cove, SK = Skye River, LN = Little Narrows, MR = Middle River.

At all sites, we caught fish by setting unbaited “Gee” brand ¼ inch mesh stainless steel minnow traps in shallow regions where we observed males courting females. These were set according to the general methods described in Schluter & McPhail (1992). Upon retrieving the traps, we evenly sampled phenotypically white and common males (identified by breeding color) and kept all females (identified by gravidity), until we had approximately 16 of each type of male and 32 unclassified females from each site. If we could not sex an individual by color or gravidity, or if a male had faded breeding colors, we did not collect it. All fish were euthanized using 0.5g/L tricaine methanesulfonate (MS-222) in seawater.

We placed all the individuals from each site into a single 1L Nalgene container containing non-denatured 95% ethanol and moved each fish to an individual 50 mL Falcon tube containing 95% ethanol a maximum of 6 hours later. Upon returning from the field (4-6 weeks after collections), we removed the pectoral and tail fins of each individual and placed the clips in 1.5mL microcentrifuge tubes filled with 95% ethanol.

### Genotyping

We extracted DNA from the clipped fins of each individual using the protocol described in Peichel et al., (2001). Briefly, the tails were digested with proteinase-K and we used a standard phenol-chloroform extraction to isolate DNA. We eluted the resultant DNA in 1X TE and assessed DNA concentrations using a Qubit flourometer (Qiagen Corp, Germany). After DNA quality control, we retained DNA from 365 individuals.

We then prepared three genotyping-by-sequencing (GBS) libraries using an adapted version of the original protocol (Elshire et al., 2011; Mondon et al., 2018). The first library contained DNA from 96 males from 2012, randomized in 96-plate well position. Based on the number of high-quality variants identified in the 2012 data, we increased the number of individuals to 148 for the second and third libraries. The two latter libraries contained DNA from a total of 296 males and females from 2014, randomized among library, plate and in 96-plate well position. We aimed for an insert size of 300-400 base pairs and used a gel-extraction method to size-select fragments from the prepared libraries. We confirmed the final fragment size distribution using a Bioanalyzer (Agilent Technologies, California). The completed libraries were then sequenced in individual lanes of an Illumina HiSeq 2000 at the University of British Columbia Biodiversity Next Gen Sequencing facility.

### Variant Identification

We identified variants using a pipeline adapted from the GATK 3.3.0 best practices guidelines (DePristo et al., 2011; McKenna et al., 2010). After demultiplexing the data using a Perl script, we used Trimmomatic version 0.32 (Bolger et al., 2014) to trim and filter sequences for quality. We then aligned the filtered reads to the stickleback reference genome v3 (Glazer et al., 2015) using BWA version 0.7.10 “mem” algorithm (Li & Durbin, 2010). We then realigned these reads using the GATK version 3.3.0 RealignTargetCreator, and IndelRealigner. Finally, we identified variants using the HaplotypeCaller and genotyped the entire dataset using GenotypeGVCFs. To facilitate analyses that required an outgroup (e.g. TREEMIX) we also identified variants from whole genome data from six marine individuals from Denmark (Ferchaud et al., 2014) and a collection of populations from the west coast (Catchen et al., 2013; Samuk et al., 2017). We processed these using the same pipeline, but with a separate run of GenotypeGVCFs.

We combined the final VCFs from the Nova Scotia and outgroup samples using the “merge” function in bcftools (Li et al., 2009). To simplify later analyses, we only included sites with biallelic single nucleotide polymorphisms (SNPs). For all analyses, we also required sites to have genotype calls in at least 80% of all individuals.

We filtered this final dataset using slightly different criteria for each analysis (described below). These filters were meant to reduce bias and/or facilitate specific statistical analyses (e.g. by reducing interdependence in the data). Unless otherwise stated, we excluded SNPs with a minor allele frequency (MAF) of < 0.05 (after Linck & Battey, 2019). For analysis requiring statistically independent sites, we pruned the dataset to reduce linkage disequilibrium (LD) between sites. This was done using the *snpgdsLDpruning*() function in the R package *SNPRelate* (Zheng, 2012). This function calculates pairwise linkage disequilbrium (*r*^2^) between all SNPs in a window of 5 SNPs, which is incremented along the genome at 5 SNP increments. If any SNPs in the window exceed the LD threshold (0.2), a single SNP is randomly chosen to be representative of SNPs in that window and the others are dropped. This reduces the statistical interdependence between SNPs caused by physical linkage, which is undesirable for most types of phylogenetic and demographic inference (Pickrell & Pritchard, 2012). The final dataset ranged from 19 000–55 000 high-quality SNPs in 354 individuals, depending on the filtration applied and the populations included.

Threespine sticklebacks have chromosomal sex determination, with males as the heterogametic sex (coded as XY, as in humans). The male sex chromosome also shares a small pseudoautosomal region with the X chromosome (White et al., 2015). Because of our reduced representation approach and poor sequencing of the Y, we chose to exclude the Y chromosome from our analysis. A more detailed analysis of the Y-chromosome of these populations using whole genome data will be presented in a forthcoming paper (Sumarli et al. 2025).

### Genetic Clustering: PCA, fastSTRUCTURE and TREEMIX

To assess whether white sticklebacks represent a distinct genotypic cluster, we first ordinated the pruned SNP data using principal components analysis (PCA) using the R package *snpRelate* (Zheng, 2012). We also examined the fit of fastSTRUCTURE models to the genetic data using the approach described in Raj et al., (2014). To further complement the clustering analyses and assess the presence of gene flow, we additionally performed tree and migration edge estimating using TREEMIX (Pickrell & Pritchard, 2012). We performed this analysis using a pruned dataset, only including individuals collected in 2014, as well as outgroup samples from Denmark and British Columbia (from Samuk et a. 2017). To assess confidence in the topology of this tree, we performed 1000 bootstrap replicates of the tree fit using the included bootstrapping mode in TREEMIX. We then combined the bootstrap trees into a consensus tree.

To test for the signal of admixture, we estimated a base tree using the Denmark samples as an outgroup, followed by sequential addition of migration edges (starting with 0). For each new migration edge, we compared the goodness of fit of the new tree to the previous tree using a likelihood ratio test (Pickrell & Pritchard, 2012). We classified the “best” tree as the last tree to offer a significant increase in likelihood. In terms of general patterns, the PCA and fastSTRUCTURE results were all highly concordant (Figure 1, Figure S2), and we thus chose to focus on the PCA results for brevity.

### Localizing the signatures of divergence

In the presence of gene flow, loci involved in reproductive isolation between two populations are predicted to exhibit unusually elevated divergence, e.g. higher F_ST_. Further, if a single large region or single locus is responsible for divergence between two populations, these extreme values should be highly localized in the genome – for example clustered in an inverted region (e.g. Küpper et al., 2016).

In order to localize the genomic regions involved in divergence between white and common sticklebacks, we performed an F_ST_ outlier analysis using the R package OUTFLANK (Whitlock & Lotterhos, 2015). This method uses a parametric bootstrap to infer the expected neutral distribution of F_ST_ (inferred via fitting a chi-squared distribution to a pruned set of SNPs) and compares the observed values of F_ST_ to this expectation to determine outlier status on a per-SNP basis. We also separately performed this analysis for the X chromosome (chromosome XIX) using SNPs from only female individuals. We did this to account for differences in (a) effective population size and (b) male-female coverage on chromosome XIX.

### Body shape

To measure body shape, we determined the coordinates of morphometric landmarks on digital photographs of individual fish using *imageJ* (Rasband, 2012). We used the landmarks described in Sharpe et al.,2008; see Figure 4B). We then imported the coordinates of these landmarks into *R* where we analyzed them using the *geomorph* 2.0 package (Sherratt et al., 2014). We performed generalized Procrustes analysis to align and scale the landmarks, followed by principal components analysis to identify the major axes of variation. We examined the first six principal components, which accounted for the majority (77%) of the variation in body shape. As found in other studies (Albert et al., 2007), the first principal component (PC1) represented differences in the degree of bending in specimens due to preservation. We thus restricted our analysis to PCs 2-5.

### Skeletal traits

To measure skeletal traits, we first stained the fish using Alizarin red following the protocol in Arnegard et al. (2014). We then took digital photographs of the stained specimens, and counted the number of lateral armor plates and then measured the length of spines using *imageJ* (Rasband, 2012). To count gill rakers, we dissected out the first gill arch and examined it under a dissecting microscope. We then counted the number of short and long gill rakers, again following the methods of (Arnegard et al., 2014).

### Body brightness

We quantified the brightness of the body by cropping a 1 cm^2^ section of the flank of each fish from a digital photograph taken under constant lighting conditions using *imageJ*. We then obtained the mean RGB values (0-255 for each channel) for these segments using *imageJ* and calculated an overall luminance score: R+G+B/(255*3).

### Testes and egg mass

We quantified testes mass by first dissecting testes from each preserved male and drying them for 36 hours in a glass desiccator containing granular desiccant. We then weighed the dried testes using a XS3DU microbalance (Mettler-Toldeo, Ohio). When both testes were developed, we weighed both and took the average of the resulting measurement; otherwise, we measured the single developed testis. We quantified egg size by extracting up to ten individual eggs from each female and weighing them in the same manner as the testes. We divided the final weight by the number of eggs measured to obtain an average egg weight for each female.

### Statistical Analysis of Morphological Data

We compared each morphological trait between white and common sticklebacks using a linear model (analysis of covariance). Because many of the traits we measured are known to covary strongly with body size, we first examined the relationship between body size (as measured by standard length or the “centroid-size” value from the Procrustes analysis, whichever was more complete) and each trait by fitting a standardized major axis regression (i.e. model II regression) using the R package *smatr* (Warton et al., 2012). If a trait displayed at least a moderately strong, significant relationship with body size (R^2^ > 0.4, p < 0.05), all further analyses were carried out via standardized major axis regression with body size as a covariate using *smatr*. Otherwise, we used standard linear models with the independent trait variable modeled as Gaussian for continuous traits or Poisson for count-based traits. When traits were measured in both sexes, we also included sex (M/F) as a covariate. For purposes of comparison between traits, we standardized all trait values by subtracting the mean and dividing by the standard deviation of the combined dataset using the R function *scale*().

An important caveat with this analysis is that the white and common sticklebacks are known to differ in body size, with whites being smaller than commons (Blouw & Hagen, 1990). This makes it difficult to statistically control fully for body size, as in some samples the two species have limited overlap in body size. In these cases, regression estimates of trait differences between species will be inexorably influenced by differences in body size, even after statistical control. The combined major axis and standard regression approach above ameliorates this issue but cannot eliminate it. As such, controlling for body size in this way will may result in an underestimate of size-independent species differences. One way to account for this in future studies would be to examine development of these traits across life stages for both whites and commons.

### Stable Isotopes

We quantified Carbon-13/Carbon-12 and Nitrogen-15/Nitrogen-14 isotopic ratios for white and common following the general method described in (Reimchen et al., 2008). Isotopic ratios of both elements have been shown to correlate with trophic position and diet in coastal marine ecosystems (France, 1995; Lerner et al., 2022). Carbon ratios can distinguish among the type of primary producer a herbivore consumes, whereas nitrogen ratios can be indicative of trophic level (France 1995, Lerner et al., 2022). We began by dissecting out the dorsal muscle from each fish, drying it in a desiccator as above, and weighing out exactly 1.00 mg of dried tissue from each individual using a microbalance. We placed each unit of weighed tissue into an individual nickel capsule. The capsules were placed in a 96-well plate, and shipped to the UC Davis Stable Isotope Facility for Carbon-13/Carbon-12 and Nitrogen-15/Nitrogen-14 ratio quantification. We did not obtain environmental samples for calibration of the isotopic ratios, and instead focused on relative isotopic differences between the whites and commons. We compared differences in Carbon and Nitrogen ratios together using a MANOVA via the *manova*() function in R (R Core Team, 2018). We included sampling location as a covariate to control for geographic variance in isotopic abundances.

## Results

### Genotypic clustering

The first two axes of our principal components analysis of the genomic data revealed three distinct genotypic clusters in the data set (Figure 1A). Males in ones of these clusters (white dots in Figure 1A) were all individuals that we had classified phenotypically as “white” in the field (light breeding colors and caught in areas with filamentous algae) in both 2012 and 2014. Two of the genotypic clusters (i.e. white and common) co-occurred at many sites across Nova Scotia (Figure 1 B &C), suggesting they do not represent neutral geographic structure. We found no clear intermediates, suggesting that our sampling did not identify any early generation hybrids, although a formal visualization of hybrid status (e.g. a triangle plot) is not possible because of the extremely low level of genetic divergence and lack of diagnostic differences between whites and commons (see later F_ST_ outlier analysis).

A third cluster (green dots, Figure 1A) consisted of stickleback exclusively from the geographically separate Bras d’Or region (Figure 1C). This cluster also contained three males that we scored phenotypically as “white” in the field, suggesting that either our field scores of phenotype were incorrect or a white male phenotype exists at a low level in the Bras d’Or population. These three individuals scored as phenotypically white did not form a separate cluster in higher PC dimensions. Note that, since that initial observation, we and others working on the white stickleback have never observed white sticklebacks in the Bras d’Or Lake region (C. Behrens, A. Dalziel and F. Chain, pers comm.).

The general results of the PCA were closely mirrored by fastSTRUCTURE (Figure S2), with the best K value being 3, determined by method of Evanno et al. (2015). Two of these three clusters again corresponded to samples with phenotypically white or common fish (males and females) from the mainland. The third distinct cluster contained all individuals sampled in the Brad d’Or Lake. At K=3, most individuals in all clusters had low levels of mixed ancestry, although there were no again no obvious F1 or backcross hybrid individuals with q≈0.5 or q≈0.25/0.75 for any ancestry source (Figure S2, K=3).

Like PCA and fastSTRUCTURE, the consensus TREEMIX tree recovered three groups of Nova Scotian stickleback (Figure 2). The white, common and Bras d’Or clades all have strong support for their individual monophyly and strong support for whites and mainland commons as relatively young sister groups (Figure 2A, 0.85-1.00 bootstrap support). The best fit for number of migration edges for the Nova Scotia/Denmark tree was three (Figure 2A, likelihood ratio test: χ^2^_1_ = 15.52, p = 0.000081). The strongest migration edge connected a western white population (Canal Lake) to another white population to the east (Sheet Harbour) (Figure 2B, red edge). The remaining two edges connected the white clade to Guysborough common populations, mirroring the signal of potential admixture from the fastSTRUCTURE analysis (Figure 2B, yellow edges).

**Figure 2.**
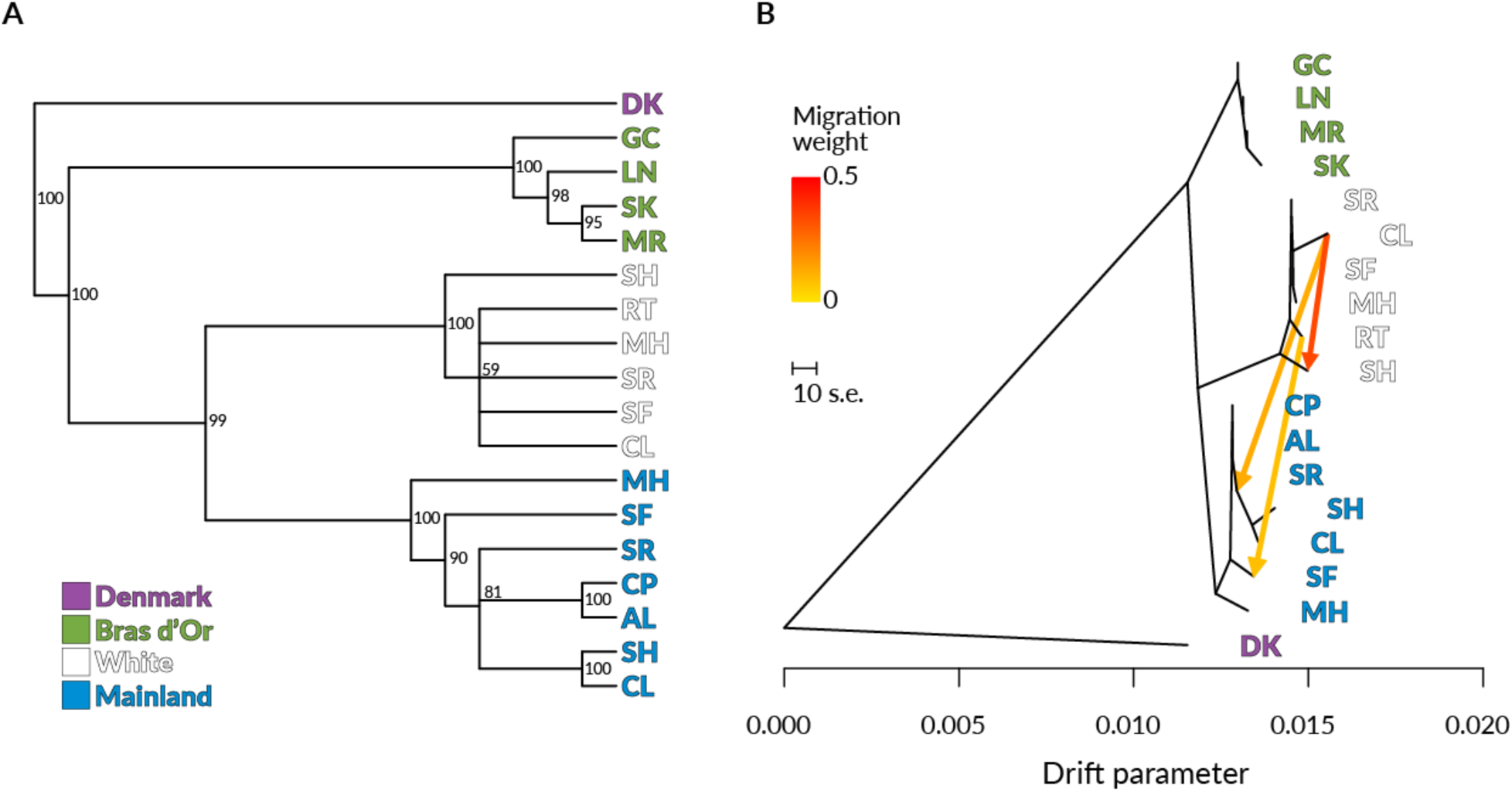
TREEMIX trees for Nova Scotian stickleback sampled in 2014, with a Denmark marine population as the outgroup. (A) A consensus tree derived from 1000 bootstrap replicates, assuming no migration. Node labels represent the percentage of trees in which each node exists. Larger numbers represent more confidence in the node. Branch lengths are arbitrary. Tip label colors correspond to genotypic clusters in Figure 1. (B) The maximum likelihood TREEMIX (m = 3 migration edges) tree. Migration edges are colored according to their weight (more red = higher relative migration). The drift parameter corresponds to the estimated amount of genetic drift that has occurred between populations.

### F_ST_ outlier analysis

Because we were primarily interested in the divergence between whites and commons per se, and not the effect of the geographic barriers to the Bras d’Or lake, we elected to restrict the F_ST_ analysis to whites and commons from the mainland only. Genetic divergence between these white and common sticklebacks occurred at many loci across the genome (Figure 3). Genome wide, we estimated Weir and Cockerham’s F_ST_ between these groups to be approximately 0.01 (95% confidence interval 0.0102-0.0110). In absolute terms, this indicates very little divergence between white and common sticklebacks: compare with the value of 0.4 between sympatric benthic and limnetic species (Schluter et al., 2025; Taylor & McPhail, 2000). Several outlier loci displayed unusually high F_ST_, with one exceeding 0.4 (Figure 3, chromosome VIII). Interestingly, these F_ST_ outliers were widespread and found at many loci across every chromosome (Figure 3). There was also no indication of the action of a single, linked block (e.g. an inversion) underlying the pattern of divergence between white and common sticklebacks. Divergence between the two types appears to be a genome-wide phenomenon, suggesting that the white and common stickleback are not simply morphs of a single polymorphic population.

**Figure 3.**
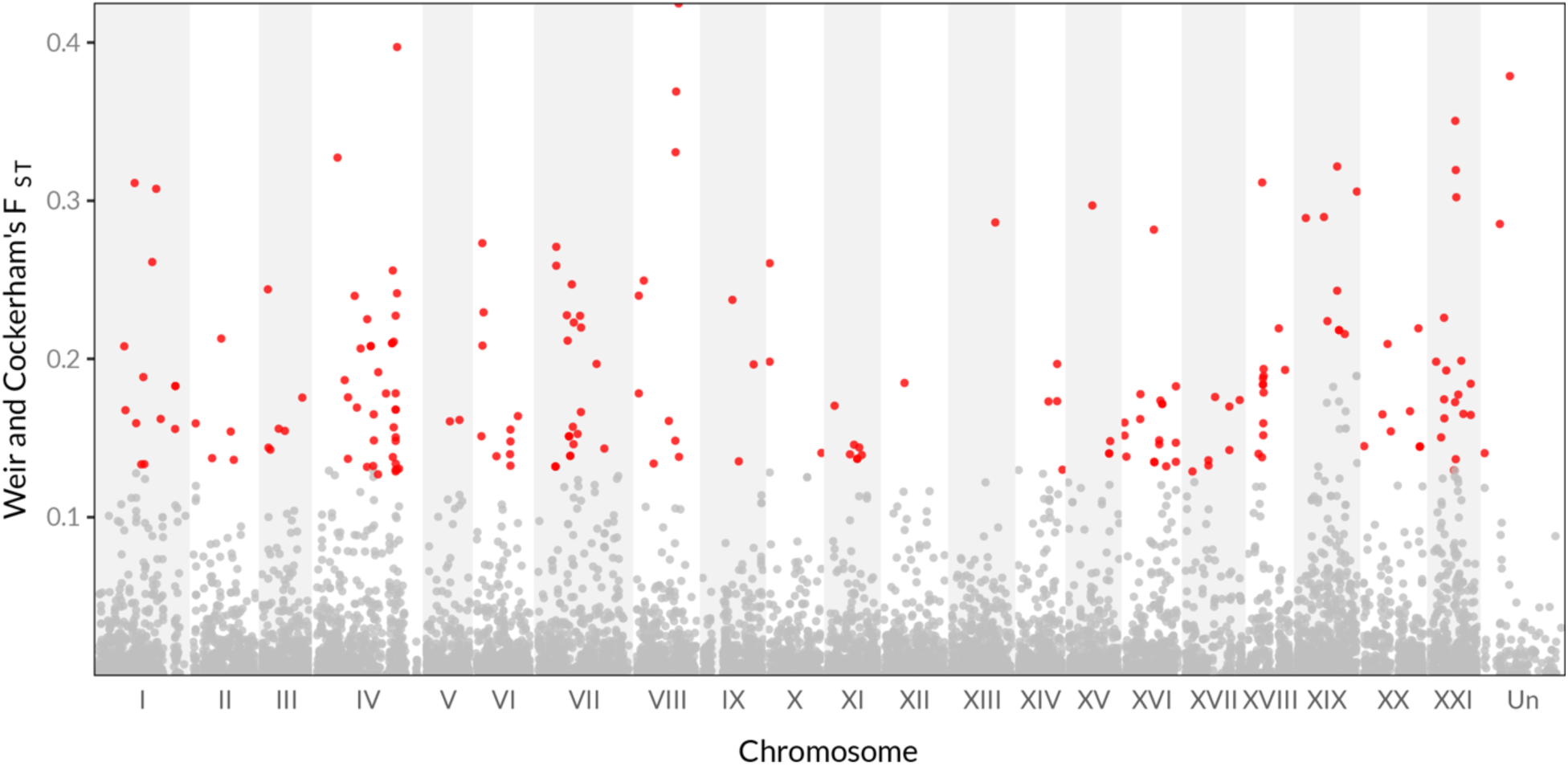
Weir and Cockerham’s F_ST_ values across the genome for the comparison of mainland white and common stickleback. Chromosomes in the reference genome (roman numerals, alternatively shaded), are plotted in genomic order with “Un” representing concatenated unplaced scaffolds (Glazer et al. 2014 assembly). Grey points represent non-outlier SNPs, and red points represent outlier SNPs identified via OUTFLANK.

### Morphological traits

White and common sticklebacks did not differ in most of the morphological characters classically associated with ecological differences in threespine stickleback, including body shape, armor plates, and gill rakers (Figure 4A). White and common stickleback nevertheless displayed moderate to large differences in a number of other morphological characters (Figure 4A). Consistent with previous work, we found that white sticklebacks are on average approximately 1.5 standard deviations (∼1.5 cm) smaller than common sticklebacks. Both male and female white sticklebacks also tend to be significantly brighter in overall color and have moderately smaller dorsal and pelvic spines than common sticklebacks, although this difference appeared to be highly sensitive to statistical control for body size (see methods). Finally, white and common sticklebacks exhibit a number of significant differences in reproduction-associated traits: white stickleback have smaller testes and eggs, but a larger number of eggs than the common form (after accounting for body size, Figure 4A).

**Figure 4.**
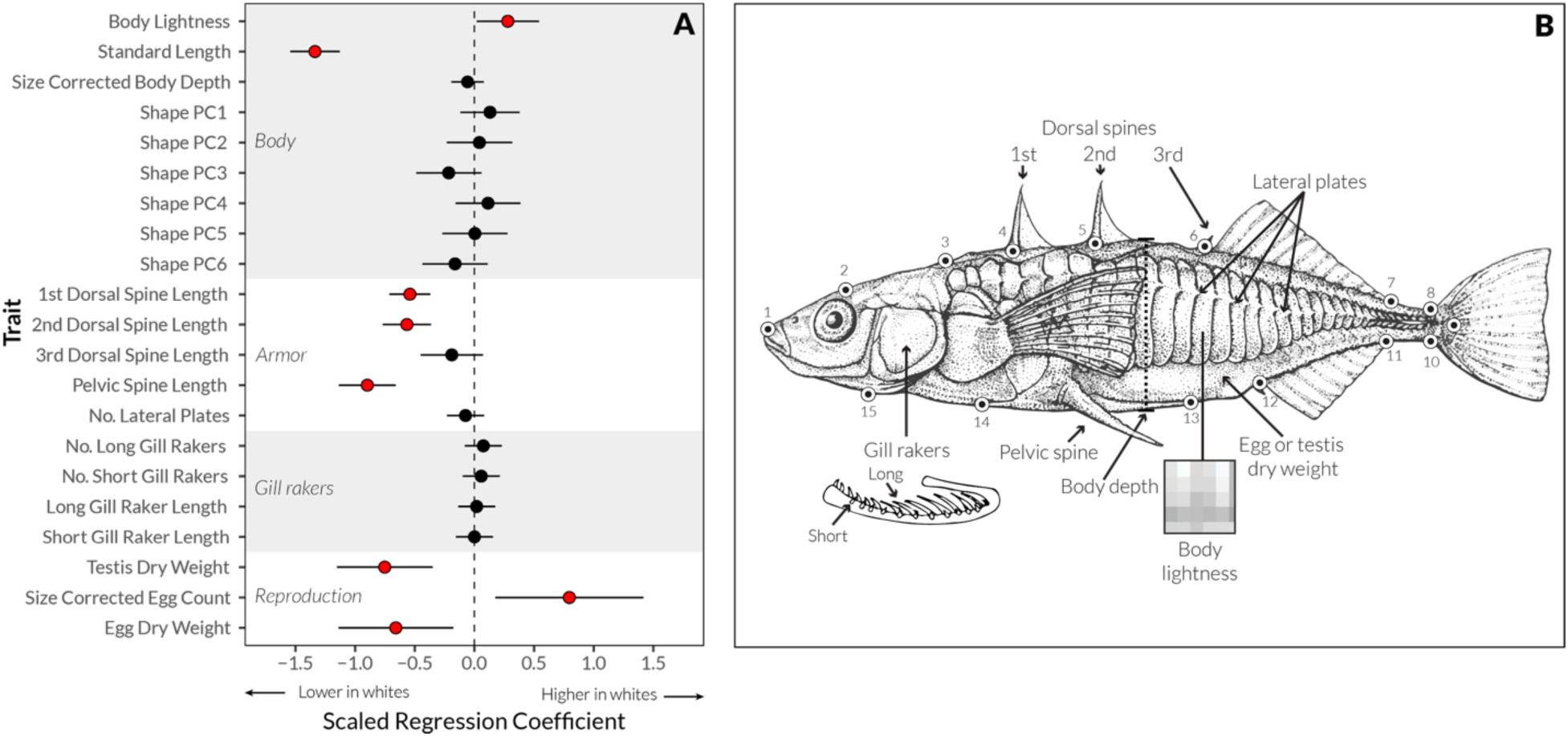
Morphological differences between white and common sticklebacks across a collection of traits with ecological and reproductive importance. (A) Model II regression coefficients of linear models fits (x-axis, units are pooled standard deviations for across all individuals) and 95% confidence intervals for differences in trait values between white and common sticklebacks. Regression coefficients include correction for the effect of both body size and sex (where applicable). “Shape PC1-6” traits correspond to scores on the first six size-corrected principal components of the morphometric landmarks shown in (B) and displayed in Figure S1. (B) A schematic of the morphological traits measured as part of the study. Artwork modified from Bell and Foster (1994) with permission.

### Stable isotopes

White and common sticklebacks do not differ in their Carbon and Nitrogen isotopic ratio profiles (Figure 5). Samples from both species show broadly overlapping distributions of both C and N ratios, and do not differ significantly in multivariate distribution (MANOVA, species term: Pillai’s Trace=0.048, Approximate F_2,106_=2.0, p=0.088). In contrast, there was a strong effect of sampling location on C and N ratios, with the Bras d’Or populations showing distinctly lower ***δ***^13^C and ***δ***^15^N ratios (Figure 5, MANOVA location term: Pillai’s Trace=1.18, Approximate F_8,214_=39, p < 2**×**10^−16^).

**Figure 5.**
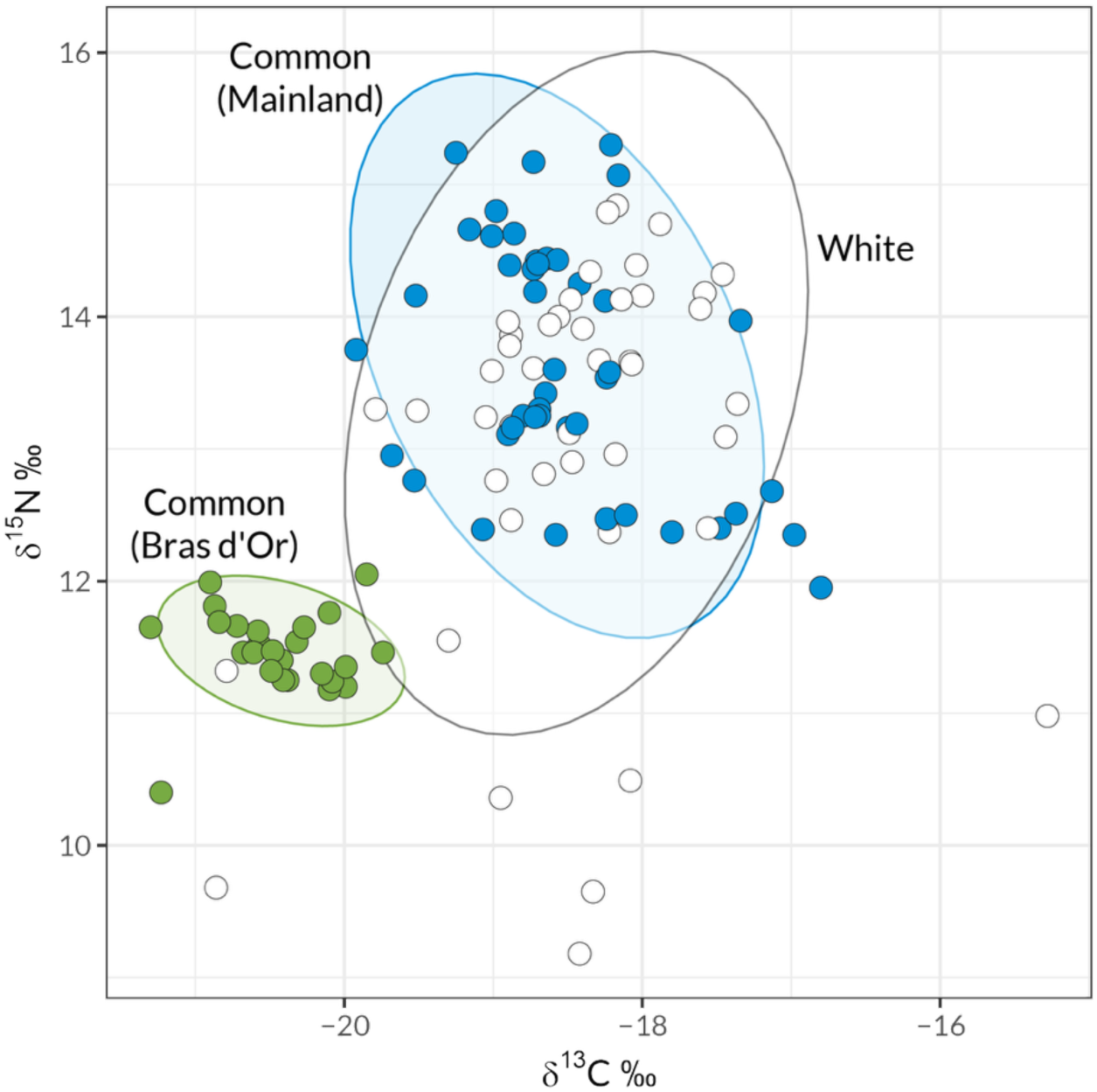
The relative abundance of ***δ***^13^C and ***δ***^15^N stable isotopes in the muscle tissue of sticklebacks collected throughout Nova Scotia. Each point represents the isotopic ratio in a single individual. Points are colored by genetic cluster as identified in Figure 1A. Shaded ellipses represent 95% confidence ellipses for reach group.

## Discussion

Incipient species in the early stages of genetic divergence provide a unique window into the causes of speciation. However, identifying such systems is inherently difficult. Here, we used modern population genomics methods and attempted to clarify whether a novel “white” threespine stickleback found in Nova Scotia is in the early stages of divergence. We found compelling evidence that the white stickleback is genetically distinct, but very closely related, to its sympatric sister species the common marine stickleback. A low level of genome-wide genetic divergence might be the result of ongoing gene flow and/or very recent divergence between the two types. We also found that white and common sticklebacks do not differ in most of the morphological characters classically associated with trophic divergence in sticklebacks except body size. The two forms were similar in body shape and gill rakers, in contrast to the sympatric limnetic and benthic species (Hatfield 1997). Their ecological similarity is reflected in their broadly overlapping stable isotope ratios. Instead, white sticklebacks appear to differ principally in male size, color and reproduction-associated traits, as well as defensive spines. As such, unlike the sympatric stickleback species pairs and all other parapatric forms such as marine-stream and lake-stream stickleback pairs (Lavin & McPhail 1993, Berner et al., 2009), the white stickleback appears to have diverged along alternative, potentially non-ecological trait axes. Instead, the white form’s most distinctive attribute is the loss of paternal care and related reproductive differences. For this reason, sexual selection and/or ecologically mediated selection on reproductive traits may be important mechanisms driving speciation in these populations. Speciation driven by these mechanisms in the absence of conspicuous ecological/trophic differences is, as far as we know, without parallel in threespine sticklebacks.

### White sticklebacks as a young species

While the population genetic methods we used indicate white stickleback are genotypically distinct from common stickleback, they are still very closely related to the sympatric common stickleback. The overall F_ST_ between white and common stickleback in Nova Scotia is only ∼0.02. For a phenotypically identifiable species, this is unusually low compared to ∼0.4 between sympatric benthics and limnetics (Taylor & McPhail, 2000, Schluter et al., 2025). It is much lower than the F_ST_ ≈ 0.2 seen between lake and stream ecotypes in British Columbia, which have relatively low levels of reproductive isolation and have not yet attained full sympatry (Roesti et al., 2012).

According to the results of the TREEMIX analysis this high genetic similarity appears to be the result of very recent divergence – most likely much more recent than the split between the Eastern and Western Atlantic 17-37 kya (Fang et al., 2018). This estimate is likely influenced by the apparent on-going gene flow between whites and commons, and higher resolution genomic data will be needed to resolve their full demographic history.

In light of the recent divergence and evidence of gene flow, it is surprising that there do not appear to be any clear early-generation hybrid individuals in our sample. For example, assuming that the genetic differences between white and common sticklebacks are numerous and found throughout the genome, hybrid individuals ought to have manifested as intermediates in the PCA projection, or as having large amounts (∼50% for an F1) mixed ancestry in the fastSTRUCTURE plots. There are several possible explanations for a lack of hybrids. First, we did not collect ambiguous-looking males, which may have been more likely to be hybrid individuals. This does not, however, account for a lack of hybrid females. It is also possible that our sampling method was somehow biased against finding hybrids. For example, perhaps hybrids have transgressive preferences for nest sites or timing of mating (such as they are in other reproductive traits, Behrens, Tucker, et al., 2025). Finally, it is possible that reproductive isolation between white and common stickleback, while recently evolved, has become nearly complete. Indeed, Blouw’s (1992) laboratory and field trials suggested that there is near-perfect assortative mating within white and common sticklebacks. If pre-mating isolation is indeed as strong as these experiments suggest, it is perhaps not surprising that we did not detect any hybrids.

### Reproductive strategy polymorphism?

The hypothesis we address here is that the white stickleback is a separate species from the common Atlantic threespine stickleback (Blouw and Hagen 1990). A key alternative explanation for the existence of a divergent white form of stickleback is that the two forms represent a male reproductive strategy polymorphism (Gross, 1996, Taborsky et al. 2008, Mank 2023). There are three types of male alternative reproductive strategy that the white stickleback could represent: genetic, ontogenetic, and condition dependent. We address the evidence for each of these below.

The white stickleback is unlikely to represent a genetically determined alternative male strategy. Alternative male strategies are predicted to have a simple genetic basis, to preclude their breakdown by recombination (Gross, 1996; Kopp & Hermisson, 2006; Taborsky et al., 2008). Empirical evidence supports this prediction, with all known cases of complex genetic mating strategy polymorphisms being associated with a large non-recombining region (Kopp & Hermisson, 2006, Küpper et al., 2016, Purcell & Brelsford 2025). The genome-wide differentiation between white and common sticklebacks is not consistent with this genetic architecture. Instead, white and common sticklebacks show genetic changes distributed over many loci and an overall reduction in gene flow despite sympatry – as expected in the case of incomplete and/or extremely recent reproductive isolation.

In contrast, a purely conditional strategy (e.g. low body condition triggers a switch in strategy) involving no genetic polymorphism would, by definition, not be strongly associated with genetic differences (Gross 1996). However, we observed that the sympatric pair of white and common sticklebacks is genetically differentiated on multiple chromosomes in both sexes at every geographic location sampled, arguing strongly against this possibility.

Finally, white sticklebacks are likely not an ontogenetically determined strategy (e.g. year one vs. year two breeding males), again because the polymorphism would not be strongly associated with genetic differences. Further, the genetic differences we see between white and common stickleback genetic clusters are stable through time (e.g. the “white” PCA cluster contains both 2012 and 2014 white sticklebacks), ruling out a cohort effect as the cause. Thus, our results are consistent with Blouw and Hagen’s (1990) original hypothesis that white and common sticklebacks represent partially (or recently completely) reproductively isolated lineages rather than alternative male strategies.

### Genetic structure within Nova Scotia

Our analyses revealed that common sticklebacks from the Bras d’Or Lake, an inland sea, are distinct from those on the outer coast (“mainland”). This is consistent with a growing appreciation for phenotypic and genetic diversity of stickleback populations in Eastern Canada, which has been generally understudied (Haines, 2023). Interestingly, in spite of previous findings by Blouw and colleagues who reported white sticklebacks in the Bras d’Or, none of the individuals sampled from Bras d’Or appeared to be genotypically similar to mainland white sticklebacks. This is made stranger by the fact that the handful of males we scored phenotypically as “white” at these populations failed to cluster with the rest of the white stickleback from the mainland. One possibility is that this is the result of the alleles that cause white nuptial colors segregating in the marine common Bras d’Or population. Perhaps persistent gene flow between Bras d’Or and whites (directly or via mainland commons) and/or balancing selection maintains a color polymorphism independent of the other loci and traits that characterize the white stickleback species on the outer coast. Further work dissecting the genetics of the white/common difference and more extensive sampling in the Bras d’Or region will hopefully elucidate these issues.

### The potential of the white stickleback study system

Our findings suggest that the white stickleback is an excellent candidate system for studying the genetics of speciation. Indeed, work has already begun to dissect the genetic basis of some of the trait differences between whites and commons (Behrens, Maciejewski, et al., 2025; Behrens, Tucker, et al., 2025). In addition, the white stickleback may serve as an excellent test-bed for theories about the interacting roles of ecological and sexual selection in speciation. White stickleback appear to have diverged in a number of mating and/or reproductive-related traits and not in most of the typical ecological traits. If this is indeed the case, sexual selection may have been an important driver of the evolution of reproductive isolation in this system. For example, white male coloration may have evolved as a sexual signal in concert with changes in nest site preferences for algae. While theory suggests that speciation via sexual selection alone is difficult in the face of gene flow (Servedio & Bürger, 2014; Servedio & Kopp, 2012), some models suggest that spatial variation in resources (e.g. nest sites) can dramatically increase the probability of speciation via sexual selection, with parental care being a potential target of selection (M’Gonigle et al., 2012; Reyes et al., 2025).

## Conclusions

Here, we used genome-wide genotyping, morphometrics and stable isotope analysis to explore whether white sticklebacks represent an ecologically-differentiated nascent species. We found that white sticklebacks are genetically distinct from common sticklebacks and that this involves multiple regions of the genome distributed over multiple chromosomes: Despite low overall genomic divergence and evidence of gene flow, male and female white sticklebacks form a unique genotypic class, distinct from sympatric common sticklebacks. As such, white sticklebacks likely do not represent a genetically determined male mating strategy polymorphism, nor young or low-condition males, but rather represent a new species with as-yet incomplete reproductive isolation. Finally, we found that white sticklebacks do not display all of the tell-tale signs of ecological differentiation found in other stickleback species pairs. Sexual selection (perhaps mediated by nest site preference) may be the key driver of reproductive isolation in this system.

## Acknowledgements

We are grateful to Max Blouw for his original work on this system, and later advice and guidance on field work and relocating his sites in Nova Scotia. Anne Dalziel, Laura Weir, and Bill Marshall all provided key support and materials when working in Nova Scotia. Kate Ostevik, Greg Owens, and Kristin Nurkowski provided essential help and guidance when preparing the GBS libraries for sequencing. Diana Rennison, Seth Rudman, and Sara Miller and members of the Schluter Lab all provided helpful edits to the original form of this paper.

## Data Availability

All code and data for the paper is available via the following Github repository: https://github.com/samuk-lab/ws_ecology

## Supplemental Material

**Figure S1.**
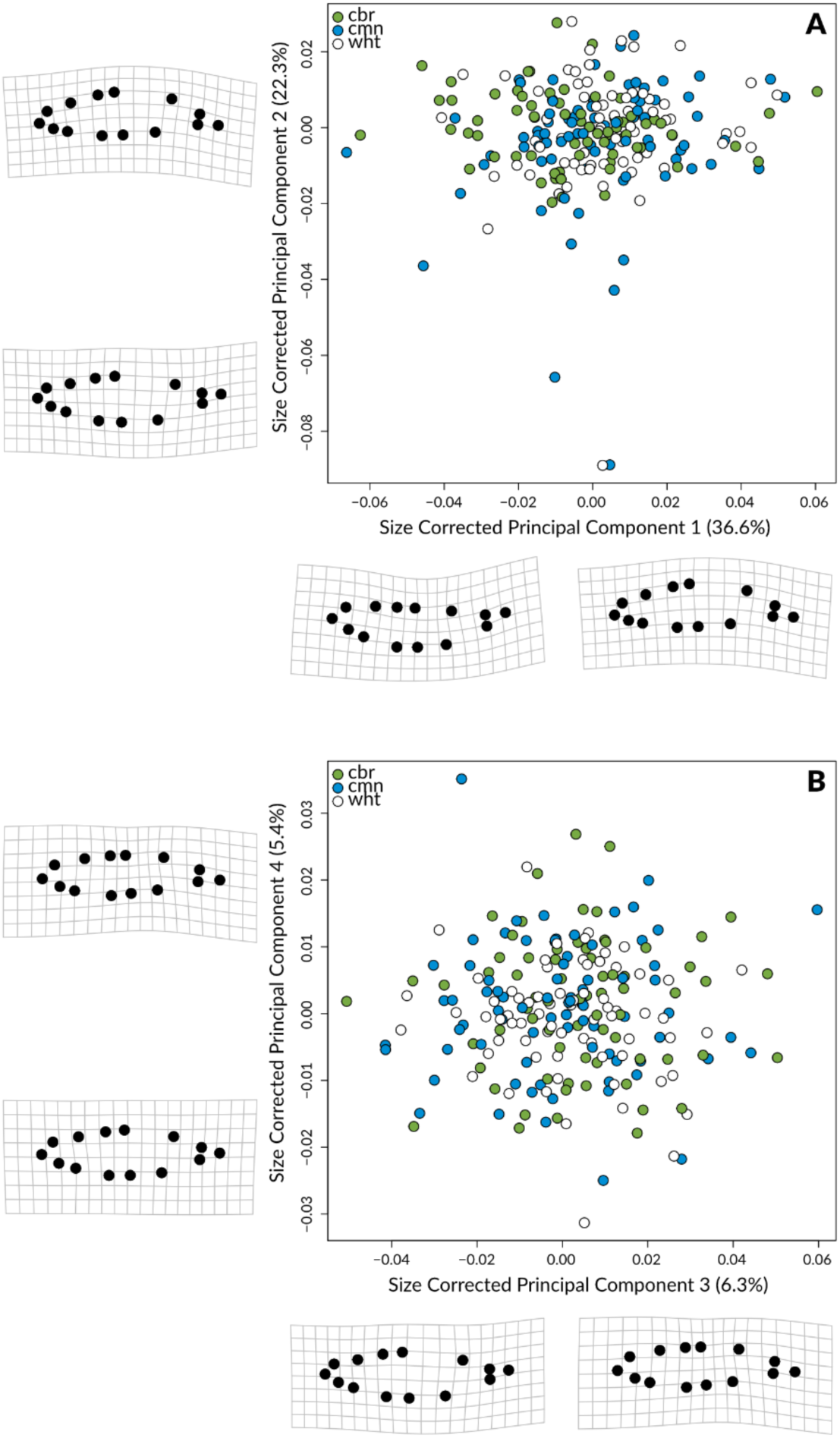
A scatterplot of a principal components scores of body shape from white (wht), common (cmn), and Cape Breton (Bras d’Or) stickleback. Dots are labelled according to their population genetic cluster assignments in Figure 1A. PCs 1-2 are shown in A, and PCs 3-4 are show in B. Thin plate spline warps and landmark positions are shown along each axis to show the range of shape variation (minimum and maximum) along each axis.

**Figure S2.**
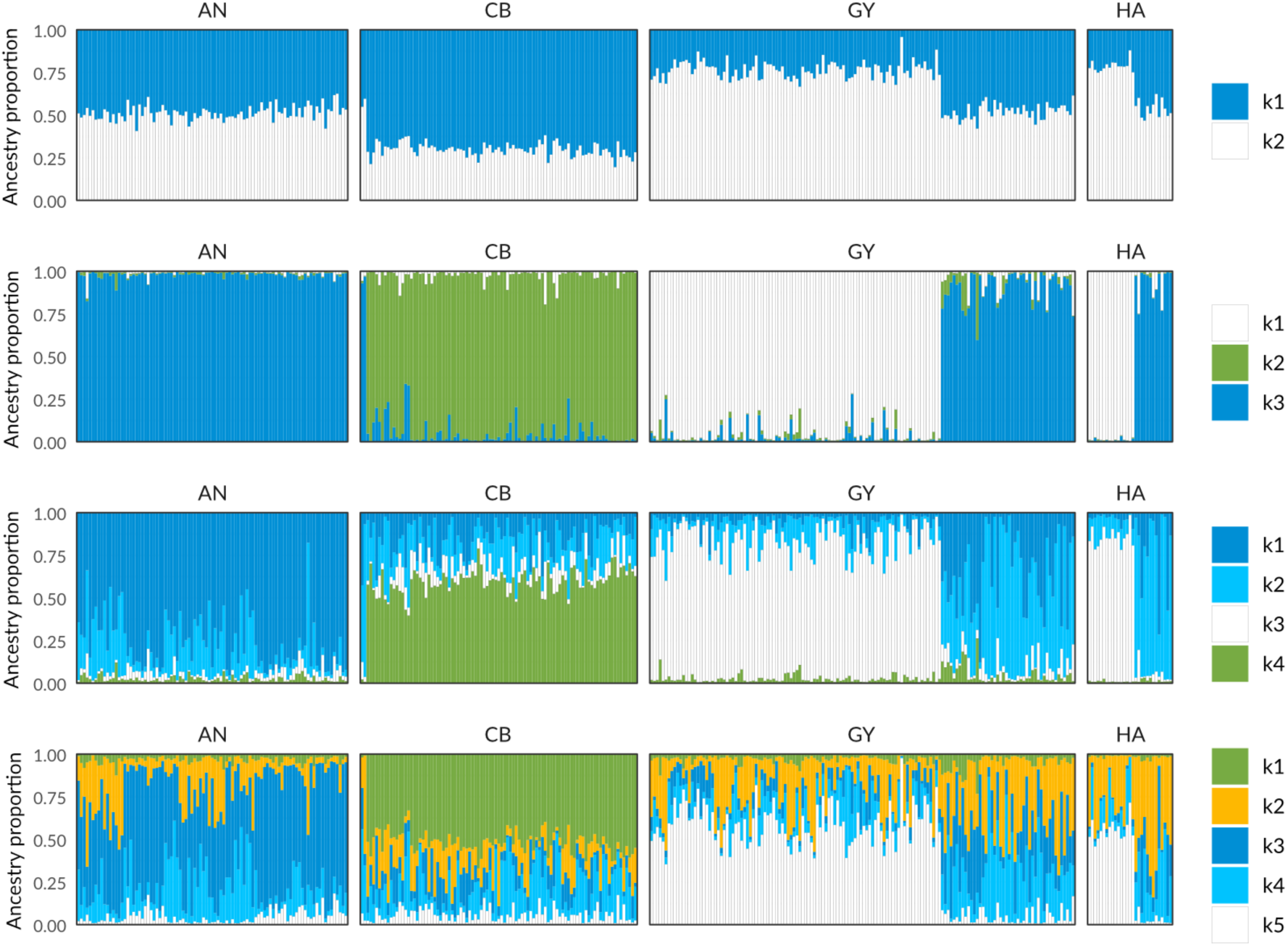
fastSTRUCTURE results for sticklebacks collected in Antigonish (AN), Cape Breton (Bras d’Or) (CB), Guysborough (GY), and Halifax (HA) regions, Nova Scotia, Canada in 2014. Each vertical bar within each subplot represents the ancestry proportions (q-value) for a single individual. Ancestry proportions are colored to match clusters in Figure 1A (main text): blue = mainland common, green = Bras d’Or common, white = white sticklebacks. Additional colors were added for clusters of unknown origin (light blue and orange in k=4 and k=5) fastSTRUCTURE results are shown for k=2 to k=5 (rows).

**Table S1.** Decimal coordinates for collection sites for white and common sticklebacks in Nova Scotia, Canada. Full site names are given in the first column along with their corresponding codes used in Figure 1A. “Types Present” refers to whether we observed the presence of both white and common types of sticklebacks or only commons (there were no sites with only white sticklebacks).

| Site Name | Types Present | Latitude | Longitude |
| --- | --- | --- | --- |
| Porper's Pond (PP) | Both | 45.43719 | -61.326 |
| Captain's Pond (CP) | Common | 45.67189 | -61.8612 |
| Pomquet (PQ) | Common | 45.62684 | -61.8448 |
| Antigonish Landing (AL) | Common | 45.63197 | -61.96 |
| Rights River (RR) | Common | 45.62721 | -61.9688 |
| Canal Lake (CL) | Both | 44.49852 | -63.9034 |
| Milford Haven Collection (MH) | Both | 45.45772 | -61.6121 |
| West Sheet Harbour Pond (SH) | Both | 44.91992 | -62.5438 |
| Black River (BR) | Common | 45.69756 | -61.0938 |
| Gillis Cove (GC) | Both | 45.91452 | -61.0545 |
| River Tillard (RT) | Both | 45.65705 | -60.9131 |
| Skye River (SK) | Common | 45.97042 | -61.1193 |
| Little Narrows (LN) | Both | 45.99216 | -60.985 |
| Middle River West (MR) | Common | 46.08322 | -60.9103 |
| Salmon River Estuary (SR) | Both | 45.35266 | -61.4728 |
| St. Francis Harbour (SF) | Both | 45.44589 | -61.3089 |

